# Digital cognitive assessments as low-burden markers for predicting future cognitive decline and tau accumulation across the Alzheimer’s spectrum

**DOI:** 10.1101/2024.05.23.595638

**Authors:** Casey R. Vanderlip, Craig E.L. Stark, Alzheimer’s Disease Neuroimaging Initiative

**Affiliations:** Department of Neurobiology and Behavior, 1424 Biological Sciences III Irvine, University of California Irvine, Irvine, CA, 92697 USA

## Abstract

Digital cognitive assessments, particularly those that can be done at home, present as low burden biomarkers for participants and patients alike, but their effectiveness in diagnosis of Alzheimer’s or predicting its trajectory is still unclear. Here, we assessed what utility or added value these digital cognitive assessments provide for identifying those at high risk for cognitive decline. We analyzed >500 ADNI participants who underwent a brief digital cognitive assessment and Aβ/tau PET scans, examining their ability to distinguish cognitive status and predict cognitive decline. Performance on the digital cognitive assessment were superior to both cortical Aβ and entorhinal tau in detecting mild cognitive impairment and future cognitive decline, with mnemonic discrimination deficits emerging as the most critical measure for predicting decline and future tau accumulation. Digital assessments are effective in identifying at-risk individuals, supporting their utility as low-burden tools for early Alzheimer’s detection and monitoring.

## 1. Introduction

Alzheimer’s disease (AD) pathologies, such as beta-amyloid (Aβ) and tau tangles, develop up to 20 years before overt cognitive decline^1,2^. Therefore, identifying individuals before cognitive decline is essential for the treatment of this disease. Significant advances have enabled quantification of Aβ and tau deposition in living individuals using both PET imaging and biofluid based techniques^3–5^. These methods are now used to identify individuals at risk for future cognitive decline and act as screening tools for clinical trials^6,7^.

While there has been substantial progress in techniques for these pathologies prior to cognitive decline, the development of sensitive cognitive tasks has lagged behind. Many of the common tasks currently used to assess cognitive function, such as the mini-mental state exam (MMSE) or clinical dementia rating (CDR), are relatively unaffected until late in disease progression ^2,8^ with impairments on these tasks lagging years behind AD biomarkers^9–11^. Therefore, it is now common to believe that cognitive decline occurs well after the buildup of pathology. While this may be the case for standard neuropsychological tests designed to measure overt cognitive impairment, there is little reason to assume that this must necessarily be the case and that pathological load could not be read out in subtle changes in cognition or behavior. If digital biomarkers can be developed and validated to reflect some aspect of AD pathology, they might offer a non-invasive, low-burden way to predict Alzheimer’s risk or monitor disease or treatment progression.

Recent work has demonstrated that digital cognitive batteries that tax circuits related to AD can accurately distinguish between individuals with cognitive impairment compared to cognitively healthy controls^12–14^. Further, longitudinal performance on these batteries is related to AD biomarkers prior to cognitive decline and are predictive of future decline on standardized cognitive tasks^15,16^. However, the tasks in these batteries are not equivalent in predicting AD biomarker status and future cognitive impairment. Specifically, tasks that tax the circuits most vulnerable to AD emerge as the best predictors of future decline.

Substantial progress has been made in understanding the neural circuits that contribute to differing cognitive functions and which circuits are particularly susceptible to AD pathologies^17–19^. Specifically, the hippocampus, a region critical for memory formation, is affected (directly and indirectly) early in the disease progression^19,20^. This vulnerability makes tasks that tax hippocampal integrity ideal candidates for detecting early cognitive changes in AD. A primary mechanism carried out in the hippocampus is pattern separation which is used to overcome competing interference between similar representations ^21–23^. To this end, it is unsurprising that one of the earliest cognitive changes in AD is the ability to differentiate between similar events ^24,25^.

Mnemonic discrimination tasks have been developed to tax hippocampal pattern separation and they show promise in identifying individuals at high risk for AD ^26,27^. Indeed, work has found that performance on these tasks is impaired in individuals with Mild Cognitive Impairment (MCI) compared to cognitively normal (CN) older adults ^28,29^. Further, these tasks can identify individuals with elevated Aβ and tau prior to cognitive decline ^30,31^. However, it is not yet fully known if mnemonic discrimination deficits are predictive of future AD pathology and cognitive decline.

Here we investigated whether subtle cognitive changes could outperform Aβ and tau at predicting future cognitive decline. We used the Cogstate Brief Battery data from ADNI as a testbed to assess the validity of digital biomarkers and demonstrate that performance on the cognitive battery better predicts conversion to MCI compared to Aβ and tau deposition. Further, we demonstrate that deficits in mnemonic discrimination drive this, suggesting that mnemonic discrimination deficits are an early marker of AD.

## 2. Methods

The data used here come from the Alzheimer’s Disease Neuroimaging Initiative (ADNI) database (adni.loni.usc.edu). The ADNI was launched in 2003 as a public-private partnership, led by Principal Investigator Michael W. Weiner, MD. The primary goal of ADNI has been to test whether serial magnetic resonance imaging (MRI), positron emission tomography (PET), other biological markers, and clinical and neuropsychological assessment can be combined to measure the progression of mild cognitive impairment (MCI) and early Alzheimer’s disease (AD). For up-to-date information, see www.adni-info.org.

### 2.1. Participants

523 older adults who took the Cogstate Brief Battery (CBB) and underwent Aβ and tau PET imaging were included from ADNI3. All participants did not have a history of major neurological or psychiatric disorders, head trauma, or history of drug abuse or dependency. Diagnosis as CN, MCI, or AD was provided by ADNI.

### 2.2. Digital Cognitive Battery

The CBB is a brief digital cognitive battery that includes four cognitive tasks, each designed to probe separate cognitive domains^32,33^. Subjects completed the battery in one sitting on a computer. All tasks involve playing cards and require “Yes” and “No” responses. The four tasks include a Detection task (DET), an Identification task (IDN), One Back Task (OBT), and the One Card Learning Task (OCL). Participants had to complete 75% of trials to be included in the study.

Descriptions of the tasks have been outlined in detail before^32,34^. Briefly, DET is a task that measures psychomotor speed where subjects click “Yes” when a playing card turns over. Psychomotor speed is calculated as the average reaction time over 35 valid trials. Invalid trials (anticipatory responses of less than 250ms) were not included in the calculations and a replacement trial was added to total 35 valid trials. IDN is a visual attention task in which either a red or black joker card flips over, and the subject responds “Yes” if the card is red and “No” if the card is black. The performance outcome of this task is average reaction time over 30 valid trials. For the OBT, individuals are shown a series of playing cards and asked if the card is the same as the previous card. This task taxes working memory and performance is quantified as average reaction time over 31 trials. OCL is a task that taxes hippocampal pattern separation which is critical for episodic memory. In this task, participants are shown a series of playing cards and are asked if they have seen the playing card previously during the task. Four cards are randomly selected to repeat eight times throughout the task. The task consists of 80 trials and the performance outcome is accuracy.

To develop a composite score reflecting overall cognitive performance, we standardized the scores from all four tasks using z-scores and inverted DET, IDN and OBT so that more negative z-scores corresponded to poorer performance. Afterward, we calculated the average of these adjusted scores across all tasks to obtain the composite measure.

### 2.3. PET Imaging

All individuals underwent Aβ PET imaging and tau PET imaging within 90 days of administration of the CBB. Individuals either underwent flobetapir (FBP) (n = 295) or florbetaben (FBB) (n = 235) imaging to quantify Aβ SUVR and flortaucepir (FTP) to quantify tau SUVR. Preprocessing of the data was handled by the ADNI PET core. Comprehensive information regarding the PET processing and acquisition techniques is available on the ADNI website at https://adni.loni.usc.edu/wp-content/uploads/2012/10/ADNI3_PET-Tech-Manual_V2.0_20161206.pdf.

Amyloid and tau PET quantification was provided by ADNI. For amyloid, a cortical composite SUVR was calculated which included the frontal, anterior/posterior cingulate, lateral parietal, and lateral temporal regions. Individuals who had a SUVR greater than 2 standard deviations above young controls (FBP: > 1.11, FBB: >1.08) were considered Aβ+ ^35^. For continuous measures, SUVR was converted to centiloids to enable comparisons across tracers^36^.

For assessing longitudinal Aβ and tau deposition, individuals with a follow-up PET scan after the initial scan were included. Both Aβ and tau SUVR annual percent change (APC) was calculated by taking the difference in uptake (centiloids for Aβ, SUVR for tau) between the initial scan and the most recent scan divided by the years between scans.

### 2.4. Statistical analyses

All analyses were done in Python and RStudio. Logistic regressions were run using statsmodels ^37^ to predict cognitive status and conversion status. Areas under the curve (AUC) measures were derived from ROC curves of the logistic regressions. Random permutation tests with 1000 permutations were used to compare AUCs between models. Commonality analyses were performed using the “yhat” package in RStudio. To identify how each variable acts in relation to the others, we performed a 6-choose-3 combinatorial analysis and quantified the number of times each metric appeared in the top third of AUCs. Pearson correlations were used to assess the associations between two continuous variables. One-way ANOVAs with Tukey’s HSD post-hoc tests were used to identify within-factor differences. For all analyses, p < 0.05 was considered reliable.

## 3. Results

### 3.1. Digital Cognitive Biomarkers perform as well and often better than Aβ and EC tau at distinguishing between CN, MCI, and AD

Significant research has shown that Aβ and tau levels are elevated in MCI and AD compared to CN older adults. Consequently, we investigated whether digital biomarkers could also differentiate between CN, MCI, and AD statuses. As expected, our findings indicate an increase in Aβ in individuals with MCI or AD compared to cognitively normal older adults with a marginal increase between MCI and AD (Fig 1A; one-way ANOVA: F(2) = 12.04, p < 0.0001, Tukey’s HSD: CN vs MCI: p < 0.0001, CN vs AD: p < 0.0001, MCI vs AD: p = 0 .09).

**Figure 1:**
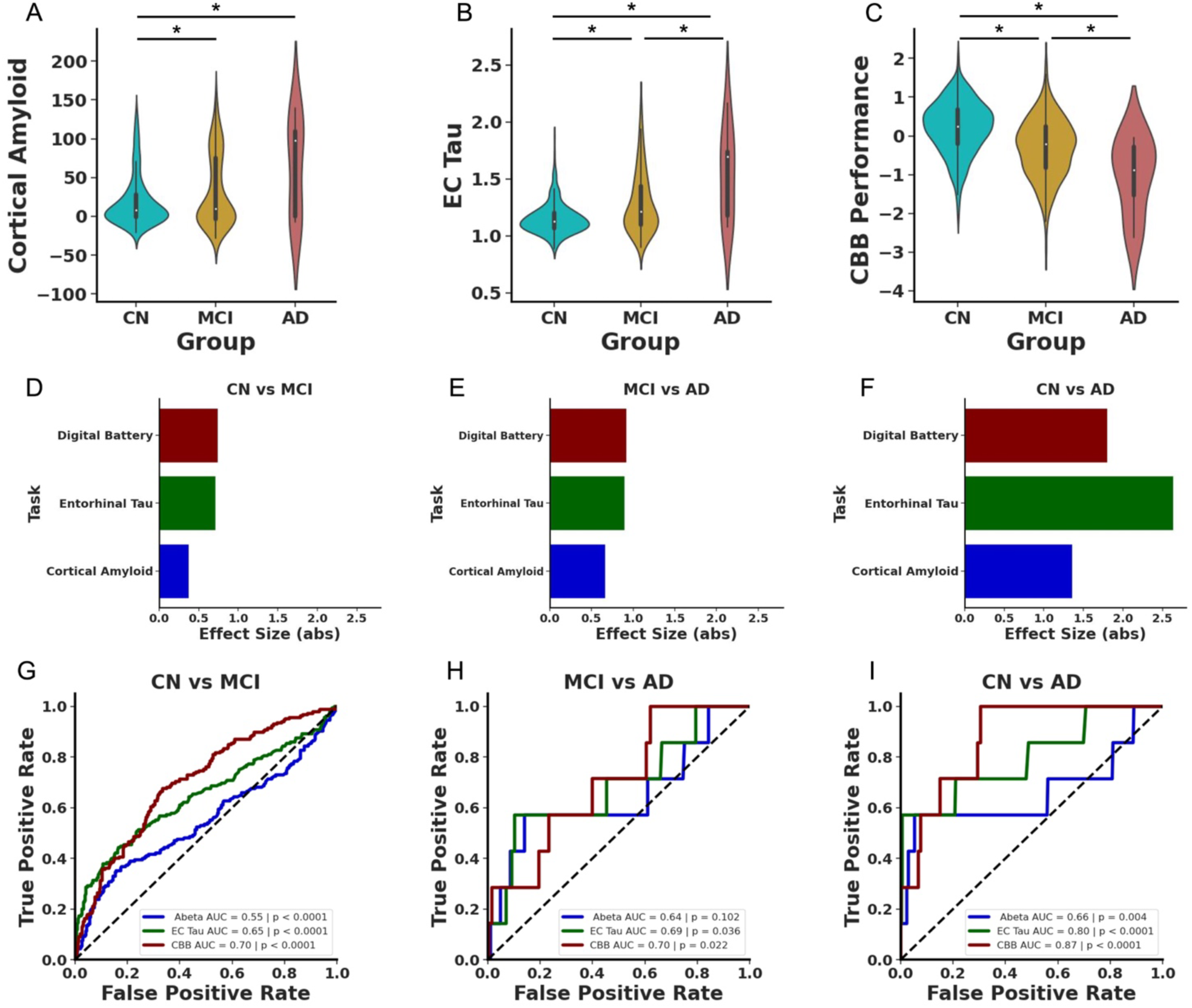
AD pathologies and digital cognitive assessments differentiate by diagnosis. A) Cortical Aβ is increased in MCI (yellow) and AD (red) compared to CN (blue) older adults. B) EC tau SUVR is increased in MCI compared to CN and further increased in AD compared to MCI. C) Performance on the digital cognitive battery declines in MCI with further impairment in AD. Effect sizes for Cortical Aβ (blue), EC tau (green) and the performance on the digital cognitive assessment (maroon) between D) CN and MCI, E) MCI and AD, and F) CN and AD. G) ROC curves show that AD pathologies and performance on the digital cognitive assessment can each differentiate CN and MCI, but the digital cognitive assessment was reliably better than the other two measures. H) ROC curves demonstrating that only the digital cognitive assessment and EC tau can differentiate MCI and AD. I) Each measure reliably differentiates CN from AD.

Similarly, EC tau was increased in MCI and further increased in AD (Fig 1B; one-way ANOVA: F(2) = 39.39, p < 0.0001, Tukey’s HSD: CN vs MCI: p < 0.0001, CN vs AD: p < 0.0001, MCI vs AD: p < 0 .01). Additionally, performance on the digital cognitive battery was related to cognitive status with CN individuals outperforming those with MCI, who in turn, outperformed those with AD (Fig 1C; one-way ANOVA: F(2) = 40.15, p < 0.0001, Tukey’s HSD: CN vs MCI: p < 0.0001, CN vs AD: p < 0.0001, MCI vs AD: p = 0.03).

Given that all measures could differentiate individuals based on cognitive task, we explored whether the degree of differences between groups varied when considering Aβ, tau, or cognitive performance. To assess this, we computed effect sizes for the differences categorized by cognitive status. We observed that cognitive performance and EC tau had roughly equivalent effect sizes for CN versus MCI and for MCI versus AD (Fig 1D) with cortical Aβ having a smaller effect size (CN vs MCI: cortical Aβ d = 0.37, CI = [0.19 0.56], EC tau d = 0.72, CI = [0.53 0.90], Digital Cognitive Battery d = 0.74, CI = [0.56 0.93]; MCI vs AD: cortical Aβ d = 0.66, CI = [−0.10 1.43], EC tau d = 0.89, CI = [0.13 1.66], Digital Cognitive Battery d = 0.92, CI = [0.16 1.69]).). Conversely, EC tau had the largest effect size for separating AD from CN (cortical Aβ d = 1.36, CI = [0.60 2.12], EC tau d = 2.64, CI = [1.86 3.41], Digital Cognitive Battery d = 1.80, CI = [1.04 2.57])

To better understand the sensitivity and specificity of these various markers, we next performed a set of logistic regression and ROC analyses using cortical Aβ, EC tau, and cognitive performance as variables to predict cognitive status (Fig 1G-I). Our analysis confirmed that each of the three measures could effectively differentiate between CN and MCI (Fig 1G). However, performance on the digital cognitive battery reached a higher AUC compared to either Aβ or tau measures (Cortical Aβ AUC = 0.55, p < 0.0001, EC tau AUC = 0.65, p < 0.0001, Digital Cognitive Battery AUC = 0.70, p < 0.0001).

We used a permutation test (1000 permutations) to determine whether any of these AUCs reliably differed. We found that performance on the digital cognitive battery and EC tau proved to be more predictive of diagnostic status than Aβ and performance on the digital cognitive battery was qualitatively better compared to tau and this approached significance (Digital Cognitive Battery vs cortical Aβ p < 0.0001, EC tau vs cortical Aβ p < 0.01, Digital Cognitive Battery vs EC tau p = 0.07). Further, all three measures could differentiate CN versus AD (Fig 1I; cortical Aβ AUC = 0.66, p < 0.001, EC tau AUC = 0.80, p < 0.0001, Digital Cognitive Battery AUC = 0.87, p < 0.0001) with cognitive performance reaching a qualitatively higher AUC compared to both Aβ and tau, but a random permutation test (n = 1000) did not find any reliable differences between AUCs (all ps > 0.15). Conversely, only EC tau and cognitive performance could reliably distinguish MCI and AD reaching similar AUCs (Fig 1H; Cortical Aβ AUC = 0.64, p = 0.10, EC tau AUC = 0.69, p = 0.04, Digital Cognitive Battery AUC = 0.70, p = 0.02) with a random permutation test (n = 1000) finding no reliable differences between models (all ps > 0.5). Thus, performance on the digital cognitive battery was at least as good as, and often reliably better than amyloid and tau at differentiating individuals based on cognitive status.

### 3.2. Deficits in mnemonic discrimination is the best predictor of MCI and AD

Given that performance on the digital cognitive battery exceeded biomarkers in detecting cognitive impairment, we next asked which cognitive domains were particularly informative by assessing performance on each of the four tasks separately. We found that performance on DET differed as a function of cognitive status with decreased psychomotor speed in MCI and AD compared to CN (Fig 2A). However, performance was not reliably different in individuals with AD compared to MCI (one-way ANOVA: F(2) = 13.99, p < 0.0001, Tukey’s HSD: CN vs MCI: p < 0.0001, CN vs AD: p < 0.01, MCI vs AD: p = 0.14). Conversely, we found that visual attention, measured via IDN, was compromised in MCI and AD compared to CN, but there was no reliable difference on IDN between MCI and AD (Fig 2B; one-way ANOVA: F(2) = 19.68, p < 0.0001, Tukey’s HSD: CN vs MCI: p < 0.0001, CN vs AD: p < 0.01, MCI vs AD: p = 0.21). Next, we found that performance on the OBT, which assesses working memory, declined in MCI and AD compared to CN, but was not reliably different between AD and MCI (Fig 2C; one-way ANOVA: F(2) = 17.92, p < 0.0001, Tukey’s HSD: CN vs MCI: p < 0.0001, CN vs AD: p < 0.01, MCI vs AD: p = 0.21). Finally, we assessed whether OCL, which measures mnemonic discrimination, declines as a function of cognitive status. We observed that performance on OCL declined in MCI compared to CN and this was exacerbated in AD (Fig 2D; one-way ANOVA: F(2) = 40.65, p < 0.0001, Tukey’s HSD: CN vs MCI: p < 0.0001, CN vs AD: p < 0.0001, MCI vs AD: p = 0.048), indicating that only OCL was sensitive to the additional decline in AD.

**Figure 2:**
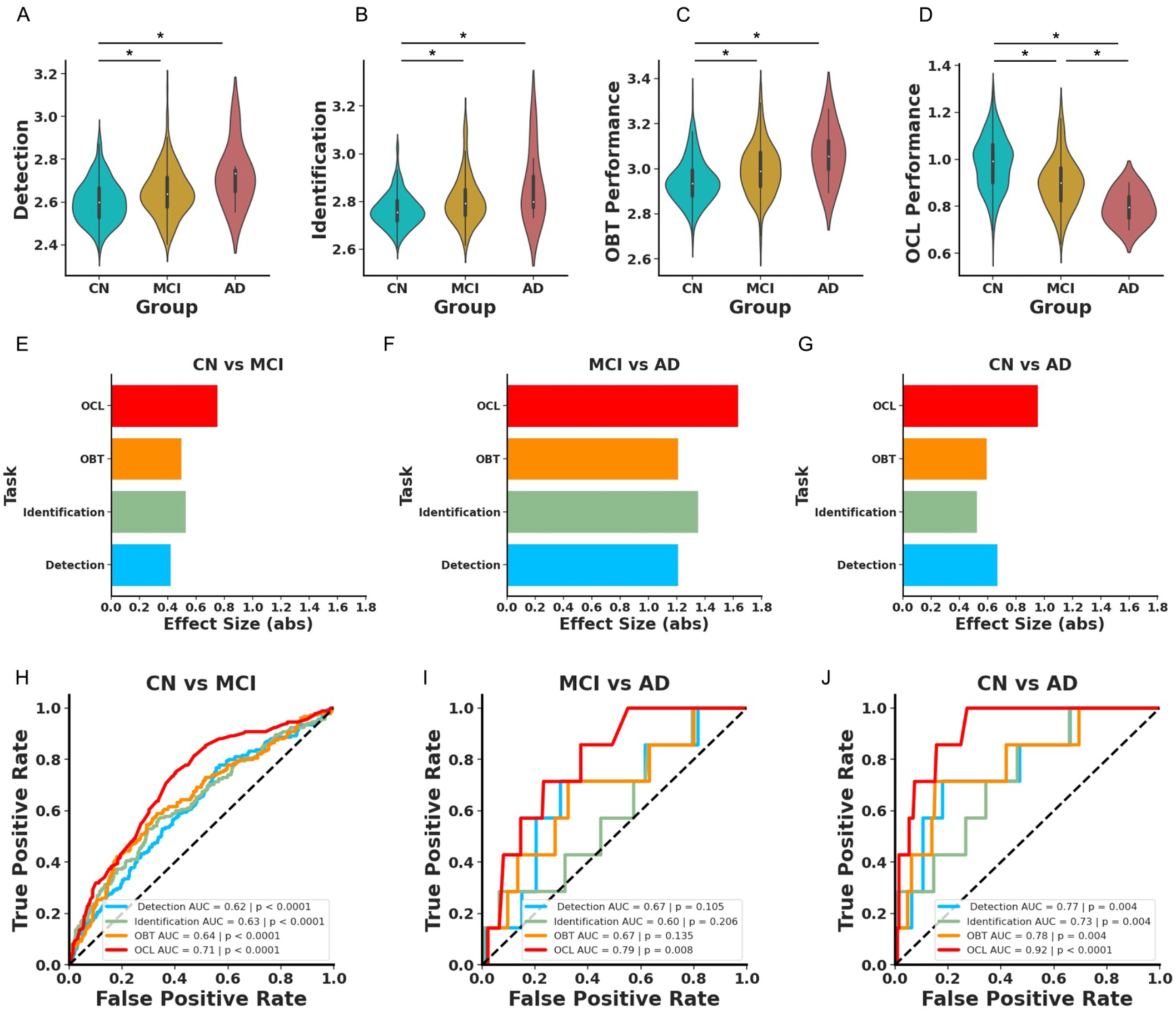
OCL is superior to other tasks for differentiating individuals by diagnosis. Violin plots depicting performance differences on A) DET, B) IDN, C) OBT, and D) OCL tasks as a function of cognitive status. Individuals with MCI (yellow) or AD (red) are impaired on all four tasks compared to CN (blue) and individuals with AD are reliably worse on OCL compared to MCI. Effect sizes for IDN (light green), DET (light blue), OBT (orange) OCL (red) between E) CN and MCI, F) MCI and AD, and G) CN and AD. H) ROC curves show that each task can reliably differentiate CN and MCI with OCL performing reliably better than the other measures. I) Only OCL performance can differentiate MCI and AD. J) Performance on each task reliably differentiates CN from AD with OCL performance reaching the highest AUC.

These findings collectively imply that MCI and AD are associated with widespread cognitive deficits. However, OCL showed the greatest difference between individuals who were CN compared to those with MCI (Fig 2E) or AD (Fig 2G) based on the magnitude of effect sizes calculated for each cognitive task (CN vs MCI: Detection d = 0.42, CI = [0.24 0.60], Identification d = 0.53, CI = [0.35 0.71], OBT d = 0.50, CI = [0.31 0.68], OCL d = 0.75, CI = [0.56 0.94], CN vs AD: Detection d = 1.21, CI = [0.45 1.97], Identification d = 1.35, CI = [0.59 2.11], OBT d = 1.21, CI = [0.45 1.97], OCL d = 1.64, CI = [0.88 2.40]). However, OCL and Detection were roughly equivalent in their effect sizes between MCI and AD (Fig 2F; Detection d = 0.67, CI = [−0.09 1.43], Identification d = 0.52, CI = [−0.24 1.29], OBT d = 0.59, CI = [−0.17 1.35], OCL d = 0.95, CI = [0.19 1.72]).

To further investigate how well these tasks separate individuals based on cognitive status, we conducted separate logistic regressions using performance on each of the cognitive tasks to predict cognitive status. The results demonstrated that all tasks could differentiate CN from MCI, with OCL reaching the highest AUC (Fig 2H; Detection AUC = 0.62, p < 0.0001, Identification AUC = 0.63, p < 0.0001, OBT AUC = 0.64, p < 0.0001, OCL AUC = 0.71, p < 0.0001). We next asked which cognitive task was the most predictive of MCI by conducting a random permutation test and found that OCL better predicted cognitive status compared to the other three tasks (OCL vs Detection p < 0.01, OCL vs Identification p = 0.01, OCL vs OBT p = 0 .03). Further, the other tasks did not vary in their predictive power (all ps > 0.28). This pattern held true when predicting CN versus AD, where again all tasks were effective (Fig 2J; Detection AUC = 0.77, p < 0.01, Identification AUC = 0.73, p < 0.01, OBT AUC = 0.78, p < 0.01, OCL AUC = 0.92, p < 0.0001) and the models did not differ in their predictive value (all ps > 0.15). However, when differentiating MCI from AD, only the OCL task showed reliable predictive capability, unlike the other tasks (Fig 2I; Detection AUC = 0.67, p = 0.11, Identification AUC = 0.60, p = 0.21, OBT AUC = 0.67, p = 0.14, OCL AUC = 0.79, p < 0.01). However, a random permutation test found no reliable differences between models (all ps > 0.16). Given that OCL better differentiated CN and MCI compared to the other tasks, we compared the predictive capacity of OCL compared to Aβ and tau. A random permutation test (n = 1000) found that OCL was superior at differentiating CN and MCI compared to both cortical Aβ and EC tau (OCL vs Cortical Aβ p < 0.0001, OCL vs EC tau p = 0.049). This suggests that OCL, which taxes hippocampal pattern separation, is particularly vulnerable to MCI and AD.

### 3.3. Deficits in mnemonic discrimination is the best predictor of progression to MCI while EC tau predicts progression to AD

We next asked whether performance on the digital cognitive assessment could better predict conversion from cognitively normal to MCI compared to Aβ and tau. To investigate this, we identified individuals who had a follow-up visit two years after administration of the digital cognitive assessment, Aβ and tau PET imaging. Individuals who were cognitively normal at baseline and remained cognitively normal two years later were called nonconverters while individuals who progressed to MCI within 2 years of baseline were called converters. We next conducted logistic regressions using either baseline digital cognitive assessment scores, cortical Aβ or EC tau to differentiate converters and nonconverters. We found that performance on the digital cognitive assessment predicted conversion over two years while Aβ and tau could not (Fig 3A; Cortical Aβ AUC = 0.57, p = 0 .08, EC tau AUC = 0.50, p = 0.24, Digital Cognitive Battery AUC = 0.74, p = 0.01). A random permutation test demonstrated that the Digital Cognitive Battery was superior to EC tau with no reliable difference between the Digital Cognitive Battery and Cortical Aβ (Digital Cognitive Battery vs EC tau p = 0.04, Digital Cognitive Battery vs Cortical Aβ p = 0.17, EC tau vs Cortical Aβ p = 0.54).

**Figure 3:**
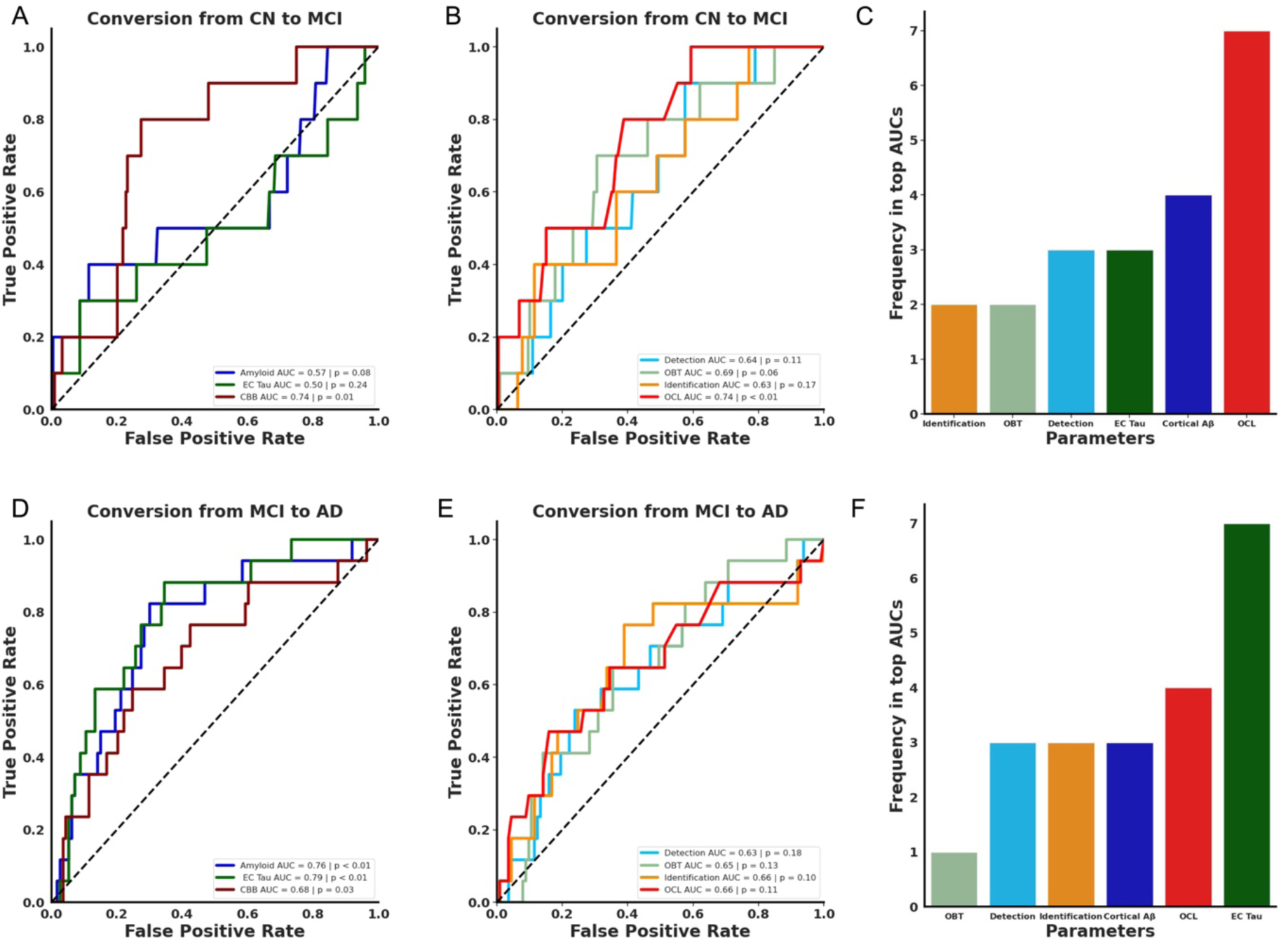
Predicting cognitive decline over two years. A) ROC curves demonstrating that performance on the digital cognitive battery (maroon) reliably predicts future cognitive decline while Cortical Aβ (blue), EC tau (green) do not. B) OCL performance reliably predicts future cognitive decline while IDN (light green), DET (light blue) and OBT (orange) do not. C) OCL appears nearly twice as often in top third of models from a permutation analysis suggesting that OCL performance is most influential in predicting future cognitive decline. D) ROC curves demonstrate that Cortical Aβ, EC tau and the digital cognitive assessment could each predict conversion from MCI to AD. E) None of the cognitive tasks could reliably predict conversion from MCI to AD. F) A permutation analysis found that EC tau was the most common metric in top third of models suggesting that this measure is important for predicting progression from MCI to AD.

To further assess whether the Digital Cognitive Battery was superior to Cortical Aβ and EC tau, we conducted a multiple logistic regression with all three measures predicting conversion status. The combined model was able to reliably predict conversion status (R^2^ = 0.10, BIC = 95.55, p = 0.04), but performance on the Digital Cognitive Battery was the only predictor that was statistically reliable after controlling for the other variables (Digital Cognitive Battery z = −2.25, p = 0.03, EC tau z = 0.45, p = 0.66, Cortical Aβ z = 1.15, p = 0.25). Further, we conducted a commonality analysis to identify which measure contributes the most to predicting conversion to MCI. We found that performance on the Digital Cognitive Battery contributed the most to the model, explaining 56.2% of the variance. Conversely, Cortical Aβ explained 18.7% and EC tau explained 1.5% of the variance (Table 1). These results suggest that the digital cognitive measures were superior to cortical Aβ and EC tau in predicting short term conversion to MCI.

**Table 1:** Commonality Analysis predicting conversion from CN to MCI.

| AD biomarkers vs Digital Biomarkers: CN to MCI |  |  |
| --- | --- | --- |
| Measure | Coefficient | % Variance Explained |
| Cortical Aβ | 0.00765 | 18.75 |
| EC tau | 0.00063 | 1.55 |
| Digital Cognitive Battery | 0.02295 | 56.25 |
| Cortical Aβ & EC tau | 0.00289 | 7.08 |
| Cortical Aβ & Digital Cognitive Battery | 0.00330 | 8.09 |
| EC tau & Digital Cognitive Battery | 0.00093 | 2.27 |
| Cortical Aβ & EC tau & Digital Cognitive Battery | 0.00245 | 6.01 |
| Digital Cognitive tasks: CN to MCI |  |  |
| Measure | Coefficient | % Variance Explained |
| OCL | 0.02195 | 54.34 |
| OBT | 0.00001 | 0.02 |
| Identification | 0.00214 | 5.30 |
| Detection | 0.00008 | 0.20 |
| OCL & OBT | 0.00003 | 0.08 |
| OCL & Identification | 0.00163 | 4.03 |
| OBT & Identification | 0.00082 | 2.04 |
| OCL & Detection | 0.00061 | 1.52 |
| OBT & Detection | 0.00001 | 0.01 |
| Identification & Detection | 0.00179 | 4.43 |
| OCL & OBT & Identification | 0.00190 | 4.71 |
| OCL & OBT & Detection | -0.00003 | -0.07 |
| OCL & Identification & Detection | 0.00355 | 8.80 |
| OBT & Identification & Detection | 0.00147 | 3.64 |
| OCL & OBT & Identification & Detection | 0.00442 | 10.95 |

To investigate which cognitive domains were the best indicators of conversion from CN to MCI, we conducted separate logistic regressions for each task. We found that only OCL could predict conversion to MCI over two years while the other tasks did not reliably predict converters (Fig 3B; Detection AUC = 0.64, p = 0.11, Identification AUC = 0.69, p = 0.06, OBT AUC = 0.63, p = 0.17, OCL AUC = 0.74, p < 0.01). A random permutation test did not find any reliable differences between models (all ps >0.32). Notably, the predictive strength of OCL was comparable to the composite score of the entire digital cognitive battery.

In a post-hoc analysis using a multiple logistic regression, we found that the overall model was modestly able to predict conversion status (R^2^ = 0.11, BIC = 100.25, p = 0.06). Within the model, OCL was the only statistically reliable predictor (OCL Z = - 2.18, p = 0.03, OBT Z= 0.08, p = 0.94, Detection Z= 0.29, p = 0.77, Identification Z= 0.42, p = 0.67) demonstrating that mnemonic discrimination is still a reliable predictor even when controlling for the other cognitive domains assessed. A commonality analysis found that OCL explained 54.3% of the variance, far more than any other task, with no other task explaining more than 5% of the variance. Importantly, only 11% of the variance was shared across all tasks suggesting that while the tasks are somewhat related, they each individually contribute to assessing the risk of progressing from CN to MCI (Table 1). To further verify that OCL was the superior measure for predicting conversion to MCI, we performed a 6-choose-3 combinatorial analysis and quantified how often each measure occurred in the top third of resulting AUCs. We found that OCL appeared in all the top models and appeared nearly twice as much as any other measure (Fig 3C). These results collectively suggest that deficits in mnemonic discrimination are predictive of conversion to MCI.

We next investigated whether these measures could predict the progression from MCI to AD over two years. To address this, we first identified individuals who were initially diagnosed with MCI and divided them into two groups: nonconverters, who remained stable with MCI, and converters, who progressed to AD. Employing logistic regressions for each measure, we found that EC tau, Cortical Aβ and the digital cognitive assessment were all effective predictors of progression (Fig 3D; Cortical Aβ AUC = 0.76, p < 0.01, EC tau AUC = 0.79, p < 0.01, Digital Cognitive Battery AUC = 0.68, p = 0.03). Further, we did not find any differences between the measures for predicting conversion to AD (all ps > 0.28).

Interestingly, in a post-hoc multiple logistic regression, we found that while the overall model was significant (R^2^ = 0.17, BIC = 102.94, p < 0.001), none of the measures could reliably predict conversion from MCI to AD (Digital Cognitive Battery Z = - 1.09, p = 0.28, Cortical Aβ Z= 1.48, p = 0.14, EC tau Z= 2.53, p = 0.11). This suggests that the measures likely share variance and therefore are not individually significant after controlling for the other measures. To this end, we conducted a commonality analysis and found that nearly half the variance (49.18%) was shared between Cortical Aβ and EC tau and 12.18% of the variance was explained by EC tau alone. Conversely, Cortical Aβ and the Digital Cognitive Battery each explained less than 10% of the variance (Table 2).

**Table 2:** Commonality Analysis predicting conversion from MCI to AD.

| AD biomarkers vs Digital Biomarkers: MCI to AD |  |  |
| --- | --- | --- |
| Measure | Coefficient | % Variance Explained |
| Cortical Aβ | 0.01313 | 9.04 |
| EC tau | 0.02456 | 16.90 |
| Digital Cognitive Battery | 0.01164 | 8.01 |
| Cortical Amyloid & EC tau | 0.07165 | 49.31 |
| Cortical Aβ & Digital Cognitive Battery | 0.00157 | 1.08 |
| EC tau & Digital Cognitive Battery | 0.00506 | 3.48 |
| Cortical Aβ & EC tau & Digital Cognitive Battery | 0.01770 | 12.18 |
| Digital Cognitive tasks: MCI to AD |  |  |
| Measure | Coefficient | % Variance Explained |
| OCL | 0.01662 | 39.30 |
| OBT | 0.00279 | 6.59 |
| Identification | 0.00259 | 6.12 |
| Detection | 0.00067 | 1.59 |
| OCL & OBT | 0.00220 | 5.20 |
| OCL & Identification | -0.00091 | -2.15 |
| OBT & Identification | 0.00371 | 8.77 |
| OCL & Detection | -0.00001 | -0.03 |
| OBT & Detection | 0.00050 | 1.19 |
| Identification & Detection | 0.00361 | 8.55 |
| OCL & OBT & Identification | 0.00036 | 0.86 |
| OCL & OBT & Detection | 0.00019 | 0.46 |
| OCL & Identification & Detection | -0.00083 | -1.97 |
| OBT & Identification & Detection | 0.01011 | 23.91 |
| OCL & OBT & Identification & Detection | 0.00068 | 1.61 |

We next asked which cognitive domains predicted conversion from MCI to AD. We conducted separate logistic regressions and found that none of the cognitive tasks could predict progression from MCI to AD (Fig 3E; Detection AUC = 0.63, p = 0.18, Identification AUC = 0.65, p = 0.13, OBT AUC = 0.66, p = 0.10, OCL AUC = 0.66, p = 0.11). These measures also did not statistically differ in predicting conversion status (all ps > 0.70). A post-hoc multiple logistic regression was not able to differentiate converters and nonconverters (R^2^ = 0.06, BIC = 119.36, p = 0.21) and none of the individual metrics were able to reliably predict converters (all ps > 0.12). In a commonality analysis, 39.20% of the variance was unique to OCL with no more than 6% of the variance being unique to any of the other tasks (Table 2). When conducting a 6-choose-3 combinatorial analysis with all the measures, we observed that EC tau was the most represented in the top third of models with OCL as a distant second (Fig 3F). This underscores the importance of EC tau in predicting progression from MCI to AD.

### 3.4. Mnemonic discrimination deficits predict future impairment on MMSE in cognitively normal older adults

Given that OCL was the most important measure for predicting conversion to MCI, we next asked whether performance on this task predicts cognitive changes in cognitively normal older adults. To assess this, we asked whether OCL performance, EC tau or Cortical Aβ predicts decline on the MMSE, a standard task used to quantify global cognitive ability. We calculated MMSE APC as the difference between the most recent MMSE score and the MMSE score at baseline divided by the difference in years. We found that baseline Cortical Aβ and EC tau were not associated with longitudinal change on the MMSE (Fig 4A; Cortical Aβ: r_p_ = −0.07. p = 0.30, Fig 4B; EC tau: r_p_ = 0.02, p = 0.76). Conversely, we found a reliable positive association between baseline OCL performance and MMSE APC (Fig 4C; r_p_ = 0.25, p < 0.0001), suggesting that impairments on OCL were related to longitudinal cognitive decline. Further, we found that there was no relationship between baseline MMSE and longitudinal changes in cortical Aβ, EC tau or OCL performance (Fig 4D; Cortical Aβ: r_p_ = 0.08, p = 0.22, Fig 4E; EC tau: r_p_ = −0.06, p = 0.44, Fig 4F; OCL: r_p_ = 0.06, p = 0.34). This suggests that OCL performance predicts future cognitive decline in cognitively normal older adults.

**Figure 4:**
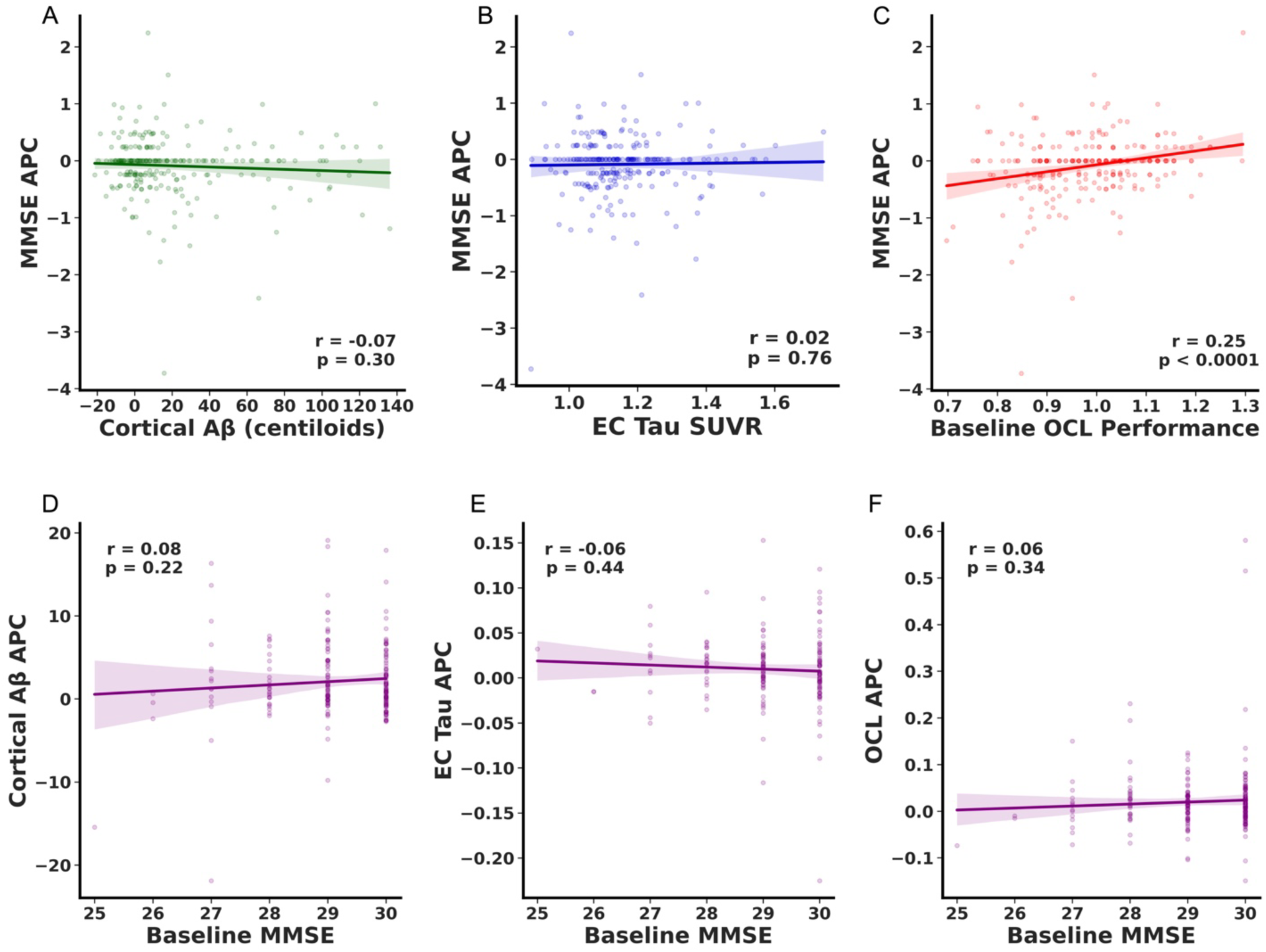
Only OCL performance predicts cognitive decline in CN older adults. No relationship between baseline A) Cortical Aβ or B) EC tau and longitudinal change on the MMSE. C) Positive correlation with OCL performance and annual change on the MMSE suggesting that lower OCL performance is associated with longitudinal decline on MMSE. No relationship between baseline MMSE and longitudinal change in D) Cortical Aβ E) EC tau or F) OCL performance.

### 3.5. Mnemonic discrimination performance predicts future tau accumulation in the entorhinal cortex and inferior temporal cortex

Given the significance of OCL performance as an indicator of future cognitive decline, we proceeded to explore whether performance could serve as a predictor for future tau accumulation in EC and IT. To do this, we correlated baseline performance on the OCL task with tau SUVR APC in Aβ- and Aβ+ individuals. Our findings revealed a significant negative correlation between baseline OCL performance and EC tau accumulation among Aβ+, but not Aβ-, cognitively normal older adults (Fig 5A; Aβ+: r_p_ = −0.26, p =0.03, Aβ-: r_p_ = −0.06, p =0.55). Conversely, there was no association between baseline OCL and EC tau SUVR APC in subjects with MCI, regardless of Aβ status (Fig 5B; Aβ+: r_p_ = −0.04, p =0.82, Aβ-: r_p_ = 0.17, p =0.33). When investigating whether OCL related to future tau deposition in IT cortex, we found a modest relationship between baseline OCL and IT tau SUVR APC in Aβ+, but not Aβ-cognitively normal individuals (Fig 5C; Aβ+: r_p_ = −0.23, p =0.07, Aβ-: r_p_ = −0.03, p =0.76). However, we did observe a significant negative association between OCL performance and IT tau SUVR APC in Aβ+ subjects with MCI, but not Aβ-MCI individuals (Fig 5D; Aβ+: r_p_ = - 0.43, p =0.01, Aβ-: r_p_ = 0.19, p =0.30).

**Figure 5:**
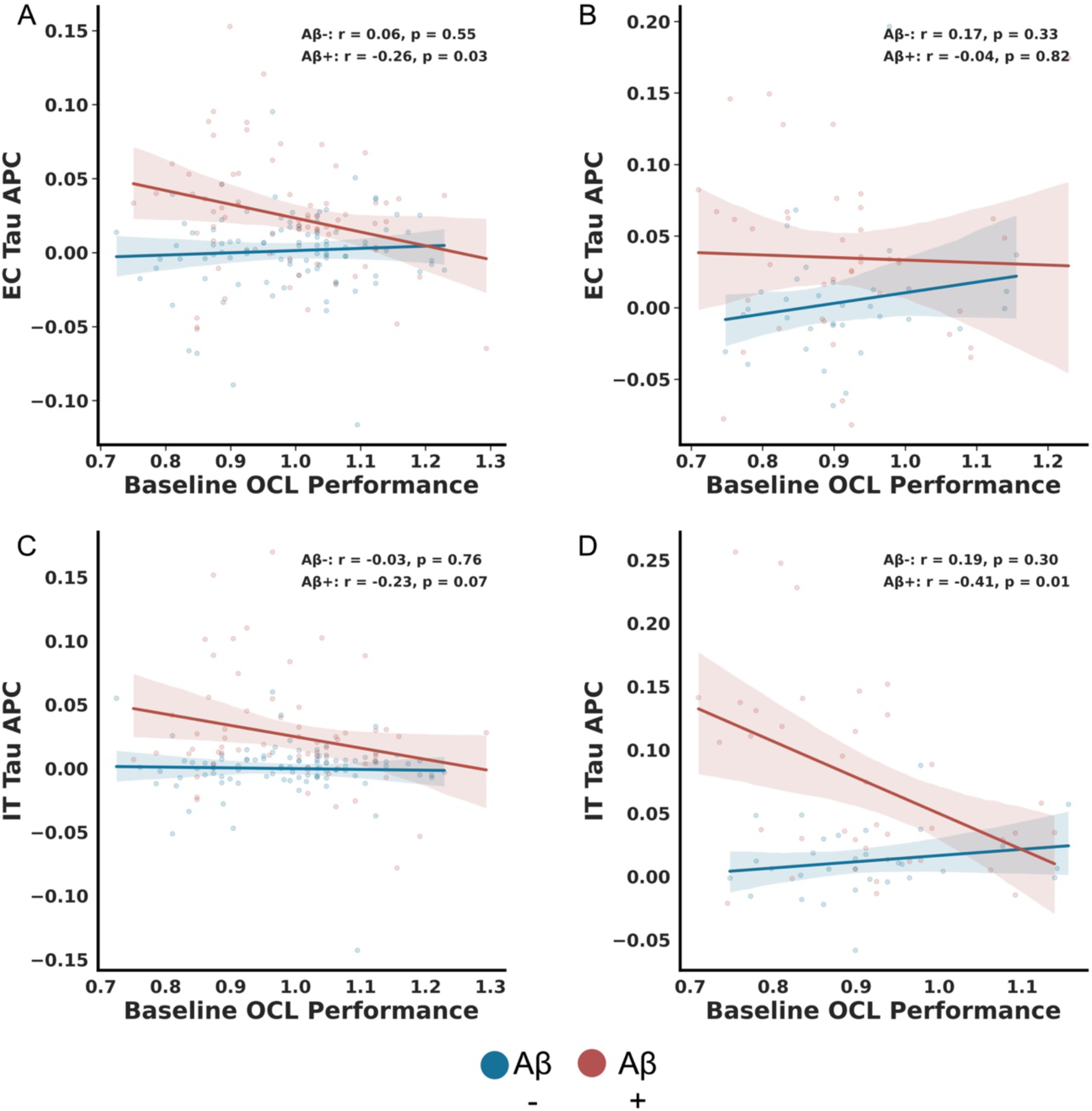
OCL is related to future tau accumulation. A) Lower OCL performance is associated with future EC tau in Aβ+ (red), but not Aβ-(blue) CN older adults. B) No reliable relationship between OCL performance and EC tau regardless of Aβ status in MCI. C) No reliable correlation between baseline OCL and IT tau accumulation in both Aβ+ or Aβ-CN older adults. D) Lower baseline OCL performance is associated with increased future IT tau accumulation in Aβ+ but not Aβ-(red) MCI older adults.

## 4. Discussion

There is a critical need for the development and validation of low burden biomarkers that identify individuals at high risk for future cognitive decline. While great strides have been made with biofluid biomarkers, less work has identified cognitive biomarkers that predict future cognitive decline. Here we used the Cogstate Brief Battery as a testbed to demonstrate that digital cognitive assessments can predict future cognitive decline. We found that the digital cognitive battery identified individuals with MCI and predicted future cognitive decline at a higher proficiency compared to baseline Aβ and tau levels. Conversely, EC tau was a critical predictor for conversion from MCI to AD. Further, we demonstrated that mnemonic discrimination deficits are the most predictive of future cognitive decline and are also related to future tau accumulation in Aβ+ older adults. This work highlights the value of digital cognitive biomarkers for identifying those at high risk for AD.

### 4.1. Utility of Digital Cognitive Batteries in identifying individuals with cognitive decline

Prior work has demonstrated the ability of digital cognitive batteries to distinguish individuals with cognitive impairment ^13,14^. Specifically, the CBB can accurately distinguish between CN and MCI at high proficiency with each of the four tasks differentiating between unimpaired and impaired older adults^38^. We replicated this in a different cohort demonstrating that all tasks can distinguish between CN and MCI. In addition, other cognitive batteries have shown promise in distinguishing between CN and MCI at high proficiency ^14,25^. However, less has been done to assess how digital cognitive assessments compare to Aβ and tau pathology in distinguishing CN and MCI. Building off these findings, we demonstrated that the digital cognitive battery was superior to both cortical amyloid and EC tau in differentiating CN from MCI, reaffirming the benefits of digital cognitive batteries.

An important question is how well biomarkers can forecast future cognitive decline. Identifying individuals at high risk for future cognitive decline can increase the therapeutic window for currently approved therapies and can aid in clinical trial recruitment. Indeed, prior work has found that Aβ and tau pathologies are predictive of future cognitive decline ^7^. However, the lack of specificity and sensitivity of these AD biomarkers suggest that other biomarkers are also needed. To determine whether digital cognitive assessments may aid in this, we asked whether performance on the battery predicted conversion to MCI over two years. We found that only digital cognitive biomarkers were predictive of future cognitive decline. Further, both a multiple regression and a commonality analysis suggested that digital cognitive biomarkers were superior to Aβ and tau levels. This suggests that digital cognitive assessments can complement Aβ and tau measures to identify those at highest risk for cognitive decline.

### 4.2. Elevated Aβ and tau is predictive of progression to dementia

The digital cognitive battery was superior to Aβ and tau for predicting progression to MCI, but we did not see the same pattern in individuals progressing from MCI to AD. In these individuals, performance on the digital cognitive assessment did predict progression, but entorhinal tau was more indicative of future decline. This aligns with prior work suggesting that tau accumulation is most rapid during MCI and relates to neurodegeneration and cognitive decline in MCI^39,40^. Importantly, in a commonality analysis, we found that nearly half of the variance was shared by cortical Aβ and entorhinal tau which suggests that these pathologies are critical for progression to dementia. Together, these results indicate that subtle cognitive changes are important for predicting progression to MCI, but once individuals exhibit overt cognitive impairment, pathologies are critical for progression to dementia.

### 4.3. Selective vulnerability of mnemonic discrimination in AD

Given that the digital cognitive battery included tasks across multiple domains, we asked whether one task was superior to the others in differentiating cognitive impairment and predicting future decline. Interestingly, we found that performance on the OCL task was most informative of cognitive status and decline. While all tasks distinguished between CN and MCI, OCL reached the highest AUC and was reliably better than the other tasks. Further, only OCL could reliably predict progression to MCI over two years. In a multiple regression model, we found that OCL was a reliable predictor of cognitive decline even when controlling for performance on the other tasks and a commonality analysis reaffirmed this, showing that OCL performance explains most of the variance in the model. Of note, MCI and AD were diagnosed cognitively, therefore, it is not completely unsurprising that OCL performance was decreased in MCI and AD. Critically, however, we compared this with other cognitive domains and Aβ and tau. Further, performance on the digital unsupervised tasks were not used in diagnosis of MCI or AD. Rather, a comprehensive in person gold standard neuropsychological testing session was used for diagnosis. Therefore, our work suggests that deficits in mnemonic discrimination were able to reliably predict impairment across the entire neuropsychological battery and at a higher proficiency than the other cognitive domains and AD biomarkers. We next asked whether OCL could predict decline on the MMSE in cognitively healthy older adults and contrasted this with Aβ and tau. We found that only OCL was related to future decline on the MMSE. A similar concern regarding cognition might apply here, but MMSE performance did not predict Aβ deposition, EC tau levels or OCL decline. This suggests that deficits on the OCL predicts global cognitive impairment, but not vice versa.

The OCL task requires individuals to remember details of playing cards despite accumulating interference and, therefore, prima facie, taxes hippocampal pattern separation. We hypothesize that hippocampal pattern separation, which reduces interference between similar representations, is particularly vulnerable to AD pathology ^27,28^. Indeed, prior work has demonstrated that performance on tasks that tax hippocampal pattern separation, such as the mnemonic similarity task, declines in MCI and individuals with AD pathologies already show impairment on these tasks prior to cognitive decline ^26,41–43^. This is likely because the hippocampus is one of the earliest areas affected (both directly and indirectly) by AD pathologies ^19^. Therefore, we propose that individuals with deficits in mnemonic discrimination are likely exhibiting declines in hippocampal integrity which is related to cognitive decline.

### 4.4. Hippocampal hyperexcitability as a predictor of future tau

Recent work has suggested that increasing tau deposition is a critical predictor of future cognitive decline ^44,45^. Specifically, it has been proposed that amyloid deposition is not pathological without tau tangles, however, amyloid can drive accumulation of tau ^46–49^. While the mechanism by which this happens is not fully understood, it’s been suggested that hippocampal hyperexcitability may be the mediating factor ^50,51^. Work has found that tasks that tax hippocampal pattern separation are vulnerable to hippocampal hyperexcitability. Specifically, increased hippocampal activity is negatively associated with performance on these tasks and pharmacologically reducing this hyperexcitability increases performance ^28,52^. Therefore, we propose that tasks that tax hippocampal pattern separation could serve as an indirect proxy for hippocampal dysfunction and in particular, hippocampal hyperactivity. This would suggest that performance on the OCL may be predictive of future tau accumulation. Indeed, we found a negative relationship between OCL performance and future EC tau accumulation in CN older adults, but only in Aβ+ individuals. Conversely, we found that OCL was related to future IT tau accumulation in Aβ+ MCI individuals. Work has found that hippocampal hyperactivity is related to future tau accumulation in both regions ^51^. However, this aligns with prior work finding that tau accumulates in EC prior to cognitive decline, but IT is particularly vulnerable later in disease progression ^11,17^. Further, the finding that this is selective to Aβ positive individuals aligns with work finding that Aβ potentiates tau accumulation. While promising, future work is needed to understand the direct connection between OCL performance and hippocampal hyperexcitability.

### 4.5. Conclusion

In this study we asked whether digital cognitive assessments could serve as low-burden biomarkers in AD. We demonstrate that performance on these assessments exceed Aβ and entorhinal tau in distinguishing CN and MCI and predicting progression to MCI. Conversely, we found that increased Aβ and tau deposition are indicative of progression from MCI to AD. Further, we demonstrate that deficits in mnemonic discrimination, which relies on hippocampal pattern separation, are informative of future cognitive decline and tau deposition. Our work suggests that digital cognitive assessments are important tools for predicting cognitive decline, and these assessments should include tasks that tax hippocampal pattern separation.

## Acknowledgements

Data collection and sharing for this project was funded by the Alzheimer’s Disease Neuroimaging Initiative (ADNI) (National Institutes of Health Grant U01 AG024904) and DOD ADNI (Department of Defense award number W81XWH-12-2-0012). ADNI is funded by the National Institute on Aging, the National Institute of Biomedical Imaging and Bioengineering, and through generous contributions from the following: AbbVie, Alzheimer’s Association; Alzheimer’s Drug Discovery Foundation; Araclon Biotech; BioClinica, Inc.; Biogen; Bristol-Myers Squibb Company; CereSpir, Inc.; Cogstate; Eisai Inc.; Elan Pharmaceuticals, Inc.; Eli Lilly and Company; EuroImmun; F. Hoffmann-La Roche Ltd and its affiliated company Genentech, Inc.; Fujirebio; GE Healthcare; IXICO Ltd.;Janssen Alzheimer Immunotherapy Research & Development, LLC.; Johnson & Johnson Pharmaceutical Research & Development LLC.; Lumosity; Lundbeck; Merck & Co., Inc.;Meso Scale Diagnostics, LLC.; NeuroRx Research; Neurotrack Technologies; Novartis Pharmaceuticals Corporation; Pfizer Inc.; Piramal Imaging; Servier; Takeda Pharmaceutical Company; and Transition Therapeutics. The Canadian Institutes of Health Research is providing funds to support ADNI clinical sites in Canada. Private sector contributions are facilitated by the Foundation for the National Institutes of Health (www.fnih.org). The grantee organization is the Northern California Institute for Research and Education, and the study is coordinated by the Alzheimer’s Therapeutic Research Institute at the University of Southern California. ADNI data are disseminated by the Laboratory for Neuro Imaging at the University of Southern California.

## Consent Statement

All human subjects provided informed consent.

